# Diversity of the dwarf palmetto: A phylogeographic analysis of the North American coastal plain and investigation of stem polymorphism in *Sabal minor* (Jacq.) Pers

**DOI:** 10.64898/2025.12.08.692648

**Authors:** Ayress D. Grinage, Jacob B. Landis, Chelsea D. Specht

## Abstract

**Premise of the study:** *Sabal minor* is considered one of two cold tolerant palm species, inhabiting riparian zones in temperate and subtropical North America. Ecologically it is often overlooked due to its short compact underground stem that makes it appear stemless, or acaulescent. However, across the central US Gulf Coast *S. minor* is polymorphic for trunk type, with some individuals identified as caulescent, bearing an above-ground trunk that can reach up to two meters in height.

**Methods:** We adopt a population genomics approach to characterize the genetic diversity of *S. minor* and to identify whether the caulescent and acaulescent forms of this temperate palm represent phenotypic variation within a single species.

**Key results:** The caulescent and acaulescent forms are not reciprocally monophyletic, suggesting these two forms are phenotypic variants of a single species, *S. minor* s.l. Genetic clustering does not correspond to stem phenotype but rather reflects five discrete biogeographic clusters that span the native range. These geographic clusters indicate phylogeographic breaks along the Tombigbee River with a history that appears to be influenced by the delta of the Mississippi River. Two ecologically divergent populations, one in White Lake and another in Central Texas, show high genetic differentiation from surrounding populations indicating the presence of ecological barriers to gene flow.

**Conclusions:** Population genetic structure of *S. minor* does not reflect stem morphotype (caulescence or acaulescence) supporting the current taxonomic recognition of one polymorphic species rather than two. However, populations of *S. minor* show genetic structuring corresponding to known phylogeographic barriers illustrating the dynamic roles regional landscape features played in shaping contemporary *S. minor*.

## INTRODUCTION

The North American Coastal Plain (NACP) is a historically underappreciated biodiversity hotspot that is highly threatened primarily due to anthropogenic habitat fragmentation (Noss et al., 2015). High biodiversity in the NACP is in part explained by the region’s role as glacial refugia during the Last Glacial Maximum (Sorrie and Weakly, 2001). Positioned south of the Laurentide ice sheet that reached as far south as Missouri, the NACP was the closet area of suitable habitat for many endemic animals and plants during glacial periods, and thus today is home to a high number of endemic species (Sorrie and Weakley, 2001). Glacial-induced migrations of widespread northern populations into suitable habitat across the NACP resulted in the division of species into isolated populations, creating opportunities for speciation and subsequent diversification. During warm interglacial periods however, connectivity between these isolated populations was re-established allowing for gene flow and/or the potential for local ecological reinforcement of species such that current population structuring may be influenced by historic periods of isolation. This temporal process of isolation followed by contact (also called expansion-contraction model) is now a well-established paradigm of glacial refugial theory and can be used to predict current population structuring of extant species (Provan and Bennett, 2008; Bagley et al., 2013). Within the NACP specifically, there is a sizable body of studies focused on identifying phylogeographic patterns of intraspecific diversity (Soltis et al., 2006) including the identification of structuring around major regional barriers e.g., the Mississippi River for insects (Stephens et al., 2011; Seal et al., 2015), reptiles (Weisrock and Janzen, 2000; Fontanella et al., 2008), gymnosperm lineages (Al-Rabab’ah and Williams, 2002), and angiosperms (Prior et al., 2020; Wang et al., 2023).

Most of these phylogeographic studies of the NACP focus on cool temperate species that migrated southward during the last glacial maximum; however, warm temperate biota of tropical origin also contribute to the biodiversity of this ecoregion. Pleistocene deposits in Louisiana contain a mixture of cool temperate (e.g., spruce) and warm temperate taxa (e.g., magnolia, tulip tree) suggesting that the US Gulf Coast was climatically mild enough for both cool and warm biota to inhabit (Deevey, 1949). Taxa like the tulip tree are implicated as relicts of older subtropical-tropical Arcto-tertiary (or boreotropical) forests because of the disjunct range between southeastern US species with closely related species in eastern Asia (Deevey, 1949). This disjunction occurred when elements of the Arcto-tertiary forests were pushed towards the tropics in response to slow global cooling that began in the Neogene and persisted through the Quaternary (Deevey, 1949; Deng et al., 2015). During glacial periods of the Pleistocene, surviving warm temperate taxa are thought to have receded into refugia in northeastern Mexico and peninsular Florida (Deevey, 1949; Martin and Harrell, 1957), resulting in a Mexican-southeastern US disjunct distribution among sister species as documented in many plant lineages (Graham, 1999) including the American sweetgum (Morris et al., 2008), American beech (Morris et al., 2010), and Black tupelo (Zhou et al., 2018). During interglacial periods populations expanded north out of Mexico and Florida and re-established connectivity between the Mexican and Florida refugia within the NACP. Therefore, warm temperate species that experienced Pleistocene glacial periods in the Mexican and Florida refugia show some degree of central Gulf Coast introgression (Martin and Harrell, 1957) in their current population-level structuring. Indeed, eastern Texas and northern Florida are considered phylogeographic “suture zones” i.e., zones containing hybrids or highly admixed populations (Remington, 1968; Swenson and Howard, 2005). Genetic signatures of phylogeographic suture zones in North America are documented in several species including the tulip tree (Sewell et al., 1996), spring peeper (Austin et al., 2002), prickly pear (Majure et al., 2012), blackstripe topminnow (Duvernell et al., 2019), and common garter snake (Jones et al., 2023). Studies documenting a northeastern Mexican and southeastern US disjunction within southeastern palms, *Sabal minor* (Goldman, 1999), and regional hybridization along the western gulf coast (Goldman et al., 2011) more broadly in *Sabal* suggests southeastern palms likely share the same phylogeographic patterning as other eastern North American subtropical species.

Palms (Angiospermae; Arecaceae) were both speciose and widespread across the northern hemisphere during the late Cretaceous and early Tertiary (Bjorholm et al., 2006) as evidenced by fossil pollen (Pierce, 1961; Harley and Baker, 2001), flowers (Allen, 2015), fruit (Manchester, 1994; Manchester et al., 2010), and leaves (Harley, 2006). Over the warm periods of the Paleocene and Eocene, palms occurred throughout middle latitude North America (Bjorholm et al., 2006; Manchester et al., 2010; Greenwood and West, 2017). Indeed, in North America, palm fossils are known as far north as Canada (Greenwood and West, 2017) and Alaska (Sunderlin et al., 2014) from the Paleocene. Late Eocene climatic cooling saw a reduction of megathermal fauna and flora at midlatitudes and dispersals south towards lower latitudes (Morley, 2003). Notably, Cano et al. (2018) identified the important role cooler climates associated with the Terminal Eocene may have played in triggering mass extinctions in two major lineages of northern hemisphere palms, the Sabaleae and Cryosophileae. For example, *Sabal* previously occurred in England (Chandler, 1978) and Germany (Mai, 1976; Manchester et al., 2010) during the Eocene, but now species are completely restricted to the Neotropics (Zona, 1990). Likewise, Grinage et al. (2024) identified the end of the mid-Miocene climatic optimum in North America as an important event that coincided with the divergence of a warm tolerant *Sabal* spp. from more cold tolerant species. Modern *Sabal* contains some of the most cold-tolerant palms (e.g., *S. minor,* Reichgelt et al., 2018) and thus, perhaps unsurprisingly, is one of the few remaining extant palm genera in eastern Northern America.

*Sabal minor* is a widespread palm species distributed from North Carolina to Central Texas (Fig. 1) occurring predominantly at the boundaries of temperate-subtropical and tropical habitats (Zona, 1990). *Sabal minor* is typically an acaulescent (“without stem” or technically bearing a subterranean stem) understory palm distributed in floodplain forests, river swamps, and natural levees (Zona, 1990). However, along the US Gulf Coast (e.g., in Texas, Louisiana, Mississippi, and Alabama), individuals of *S. minor* deviate from their acaulescent form to produce populations with individuals exhibiting both the acaulescent type as well as a continuum of caulescent forms where individual palms have been documented with stems as tall as two meters (Fig. 1). These caulescent growth forms were historically considered a distinct species from *S. minor,* described as *S. louisiana* (Darby, 1817; Small, 1926, 1929; Bailey, 1934, 1944; Bomhard, 1935, 1943; Ramp and Thien, 1995). Despite being described as a separate species, there was an overall paucity of morphological, ecological, and enzymatic data to suggest that the two growth forms functioned as distinct species (Bailey, 1944; Zona, 1990; Ramp and Thien, 1995).

**Figure 1:**
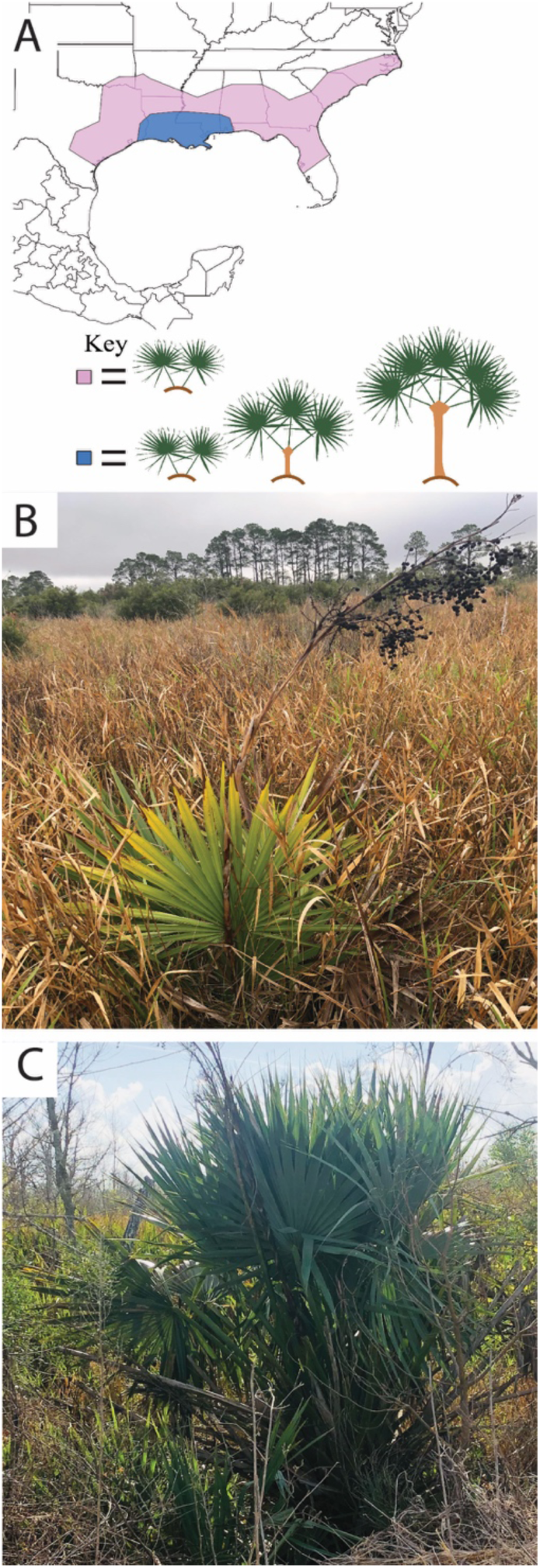
Growth forms of *Sabal minor*. Geographic distribution of *S. minor*. (A) Map of the natural distribution of *S. minor* (Zona 1990). The pink region denotes where only the acaulescent form grows, while the polymorphic region where both the acaulescent and caulescent growth forms occur is shown in blue. (B) Acaulescent growth form and (C) the caulescent form from White Lake Conservation Area, Gueydan, Louisiana.

*Sabal minor* is currently the most northernly distributed palm in North America. Based on recent studies, *Sabal minor* was likely distributed across the NACP as recently as 12 million years ago (Grinage et al., 2024). Considering the possible age of *S. minor* and its distribution, *S. minor* likely experienced historical episodes of climatic warming and cooling, and genetic footprints of these geologic periods are present across the evolutionary history of the species. We expect that the observed phylogeographic patterns reflect the physiographic history of the region particularly along fresh waterways due to the distribution and ecology of *S. minor*. We also suspect we will observe some level of phylogeographic or ecological patterning according to stem morphology (Fig. 1) similar to the results of Ramp and Thien (1995). A key motivation of this work is resolving whether the acaulescent and caulescent growth forms are separate species or are variants within *S. minor*. In this study, we use population genetic tools to assess population-level clustering and genetic admixture to assess patterns of genetic diversity across the distribution of *S. minor*. We use these analyses together with a phylogeographic analysis and inferences of migration events to evaluate broader geographic and ecological processes structuring the genetic diversity of *S. minor*.

## METHODS

### Assessment of *S. minor* phylogeography

#### RAD-seq

A total of 95 samples were collected for this study (Table S1). Of these, 79 accessions are sampled from across the native range of *S. minor* in the southeastern US (Table S1) representing individuals of the caulescent (n= 27) or acaulescent (n= 45) habit; seven individuals have an unknown status due to lack of data from sampled herbarium specimens. Specimens represent known polymorphic populations (Fig. 1) within Alabama, Mississippi, Louisiana, and Texas with additional populations from North Carolina, South Carolina, Georgia, and Florida. Most of the sampled habitats consist of floodplain, riparian forests or hardwood hammocks; however, one locality from Gueydan, Louisiana (White Lake Conservation Area) is freshwater marshlands (Fig. 1B, C). All samples identified as *S. minor* (acaulescent or caulescent) were characterized by their minimally intruded costa into the leaf blade which results in a flat-shaped blade as opposed to a curved blade (c.f. *S. palmetto*), and the presence of an erect inflorescence (Fig. 1B, C). One horticultural variety of *S. minor*, the “Emerald Isle Giant”, which is a large, acaulescent specimen cultivated from the personal garden of Dr. Kyle Brown with provenance from Emerald Island, North Carolina was also included. Sixteen samples were included as outgroups: *Sabal x brazoriensis* (n= 2), *S. tamaulipensis* (n= 3), *S. palmetto* (n= 5), *S. miamiensis* (n= 3), *S. etonia* (n= 2), and *S. mexicana* (n= 1). All accessions collected from living material (either wild or cultivated) were preserved directly on silica gel and stored at room temperature until DNA was extracted.

#### DNA Isolation and Sequencing for RAD-seq

High quality DNA was extracted from leaf tissue (either silica dried or from herbarium specimens) following a modified CTAB protocol (Doyle and Doyle, 1987), see genome details in Appendix S1 for more extraction details. Extracted DNA was cleaned using homemade Ampure beads (Rowan et al., 2017) following the same protocol as De La Cerda et al., (2023). Cleaned DNA was run on a 1% agarose gel (up to 1 hour at 100 V) to evaluate DNA degradation and the presence of high molecular weight DNA. High quantity DNA was identified as that having a qubit concentration of 40 ng/µL or more. The concentration of DNA used for library preparation was between 20-60 ng/µL in 30 µL (except for three accessions that were 11.6, 13.2, and 80.2 ng/µL). Extracted DNA was sent to the University of Wisconsin Biotechnology center for library preparation. Libraries were prepared for paired-end RAD sequencing (RAD-seq) using the *apeK1* enzyme. Samples were pooled and sequenced on one lane of Illumina HiSeq 4000 2×150 bp at Novogene (Sacramento, CA).

#### Reference mapping and SNP filtering

To aid SNP assembly, we generated a hybrid genome assembly using short- (Illumina) and long-read (Oxford Nanopore) data for one accession of *S. minor* (Appendix S1). This sequence data was used to generate a genome assembly using MaSuRCA v4.0.7 (Zimin et al., 2013). Following assembly and polishing, we used the resulting hybrid genome assembly as a reference to map raw reads and call SNPs. Raw rad-seq reads were demultiplexed with the process_radtags script in STACKS v2.62 (Catchen et al., 2013).

Demultiplexed reads were then aligned to the assembled reference genome using BWA-mem2 v2.2.1 (Vasimuddin et al., 2019). Aligned reads were used to identify single nucleotide polymorphisms (SNPs) using the ref_map.pl script within STACKS. One accession of *S. minor*, (SMPGL59) was excluded as it only contained 224,004 raw reads, well below the average (9,375,982 reads). For the remaining 94 samples, VCFtools v0.1.13 (Danecek et al., 2011) was used for data filtering. SNPs were filtered to retain only biallelic sites (--min-alleles 2 –max-alleles 2), remove sites with low minor allele frequency (--maf 0.05), low minor allele count (--mac 3), and more than 50% missing data (--max-missing 0.5). To leverage SNP data to infer biological processes, SNPs need to represent freely recombining (unlinked) loci (O’Leary et al., 2018). Thus, further filtering based on linkage disequilibrium was done after estimating linkage disequilibrium (LD) with PopLDDecay v 3.30 (Zhang et al., 2019) to identify the LD correlation coefficient (r^2^) for a sliding window trimming approach. We used the lowest r^2^ threshold (r^2^= 0.4), where the LD decay plot started to become stationary (Fig. S1), an indep-pairwise approach in PLINK v1.90b6.21 (Purcell et al., 2007), with a sliding window size of 20 kb and step size of 10 kb.

Given that ad hoc filtering for dataset features such as missing data can bias results (O’Leary et al., 2018; Linck and Battey, 2019), seven additional datasets were generated to evaluate the effect of filtering on the results of a principal components analysis (PCA) in differentiating clusters of all 94 individuals. We tested filtering regimes under varying thresholds of Minor Allele Frequency (MAF): 0.02, 0.04, 0.06, 0.08, 0.10 and Minor Allele Count (MAC) = 3, max-missing = 0.5. We also evaluated the effect of missing data (--max-missing 0.9) and (--max-missing 0.7); MAC = 3, MAF = 0.05. Linkage pruning was performed on all seven datasets as described above, followed by PCA (see details below). With varying levels of MAF and missing data allowed, we saw evidence of data distortion due to filtering when MAF = 0.02, 0.08, 0.10 in PC2 vs PC3 and PC3 vs PC4, as shown by varied genetic clusters of species in the PCA (Fig. S1). Although we did not see evidence of data distortion when MAF = 0.04-0.06 and with only 30% and 10% missing data allowed, for all downstream analyses we used the filtered data set with MAF = 0.05 and 50% missing data allowed to maximize SNP retention while balancing for SNP filtering.

#### Phylogeographic Relationships and Historical Migration

Phylogeographic relationships were inferred using 92 specimens: 78 *S. minor* individuals and individuals from 12 other *Sabal* species. The two hybrid accessions, individuals of *S. x brazoriensis,* were excluded from analyses that assume strict bifurcating relationships. The python script VCF2Phylip (Ortiz, 2019) was used to convert the filtered and LD-pruned vcf file for all 92 accessions to a fasta file. We ran RAxML-ng v1.2.0 (Stamatakis, 2014) to obtain a maximum likelihood (ML) tree under the GTGTR+G4 model the substitution model based on Bayesian Criterion Information (BIC) score in ModelTest-ng v.0.1.7 (Darriba et al., 2020). To assess support, we implemented 1000 bootstrap pseudoreplicates. The bipartition tree was visualized in FigTree v1.4.4 (Rambaut, 2009). To assess conflicting signals within the data, we ran SplitsTree v6.0.29-beta (Huson and Bryant, 2006) and SVDquartets (Swofford, 2003). We used the full data set including the two *S. x brazoriensis* accessions (94 accessions) in SplitsTree.

#### Calculating Genetic Diversity and Differentiation

We used a principal components analysis (PCA) to visualize genetic clustering of 78 accessions of *S. minor* and 16 outgroups using the R package SNPRelate v1.34.1 (Zheng et al., 2012). Genetic structure and observed admixture were assessed with the LEA v3.12.2 (Frichot and François, 2015) package in R on only *S. minor*. To calculate the best K value indicating potential number of ancestral populations K= 1-25 with 200 iterations for each were evaluated. The best fit K was validated under a cross-entropy criterion where the best fit K was selected at the lowest trough (Fig. S3). Genetic differentiation for each locality and subsequent populations identified by the PCA were calculated in the R package SNPRelate using pairwise population weighted Fst.

To evaluate population-level admixture and migration, we used TreeMix v1.13 (Pickrell and Pritchard, 2012) which models migration events and admixture based on a ML tree. Eight populations were used to bin the 78 specimens of *S. minor* into five populations based on the results from LEA and SNPRelate, with three outgroup species: *Sabal palmetto* (n=5 accessions), *Sabal tamaulipensis* (n=3 accessions), and *Sabal mexicana* (n=1 accession). Additionally, we used Adegenet v.2.1.10 (Jombart, 2008) to interpret spatial genetic differentiation for isolation by distance (IBD) with a mantel test consisting of 10,000 simulations. We estimated IBD with different groupings of the samples (1) across all *S. minor* (n=78), (2) across major modern river basins: Atlantic Coast (n= 13), Central Gulf Coast and White Lake (n= 57), Central Texas and West Gulf Coast (n= 8), and (3) across populations on each side of the Mississippi River, east: Atlantic Coast and Central Gulf Coast (n= 64) and west: White Lake, West Gulf Coast, and Central Texas (n= 14) of the Mississippi. Population statistics including number of private alleles, nucleotide diversity (p), and inbreeding coefficient (FIS) were calculated using only variant sites in STACKS (--fstats).

## RESULTS

### Genome Assembly

The hybrid MaSuRCA genome assembly (CoGe accession ##) resulted in an assembly of 703 MB in 2679 contigs with an N50 of 745,027 bp. The BUSCO analysis showed 97.9% complete BUSCO genes (1580 total with 1458 complete and single copy and 122 complete and duplicated), 1.2% fragmented (20 BUSCO genes), and 0.9% missing (14 genes) out of the total of 1614 genes in the embryophyta version 10 database.

### RAD-seq data

Approximately 44 GB of raw data were generated from the paired-end libraries. One sample (SMPGL59) was poorly sequenced and thus excluded before mapping to the reference. Mapping raw reads to the genome and calling SNPs yielded 1,324,320 loci and 5,148,690 SNPs prior to filtering. The average percent missing data per individual was 23.3%; with the herbarium specimens having 33.9% missing data and the silica specimens 22.3% (Fig. S4).

### Phylogeographic Relationships

Both RaxML and SVDquartets indicated that *S. minor* forms a well-supported clade (Fig. 2 and Fig. S5). Within *S. minor,* accessions from caulescent and acaulescent individuals do not form unique or reciprocally monophyletic (Fig. 2). Rather, clades were resolved corresponding to the geographic location of the individual accessions. Accessions from the Atlantic Coast (Fig. 2, purple) form a grade at the base of the *S. minor* accessions but appear as a clade sister to all other accessions in the SVDquartets phylogeny (Fig. S5). A clade with individuals from White Lake (orange) is nested within a clade of individuals from the Central Gulf Coast (blue). Individuals from the West Gulf Coast (pink) and Central Texas (green) together form a clade that is also nested within a clade of individuals from the Central Gulf Coast (Fig. 2). The Central Gulf Coast accessions themselves form a large clade that can be divided into three subclades, that give rise to the White Lake, the West Gulf Coast, and Central Texas (Fig. 2). SplitsTree showed the same regional aggregation as both phylogenies (Fig. S6). SplitsTree showed an alternative placement of one accession of *S. x brazoriensis* (SMPGSbraz2_C2) which was nested within *S. minor* from the West Gulf Coast (Fig. S6) indicating a potential hybrid origin with a member from this West Gulf Coast population.

**Figure 2:**
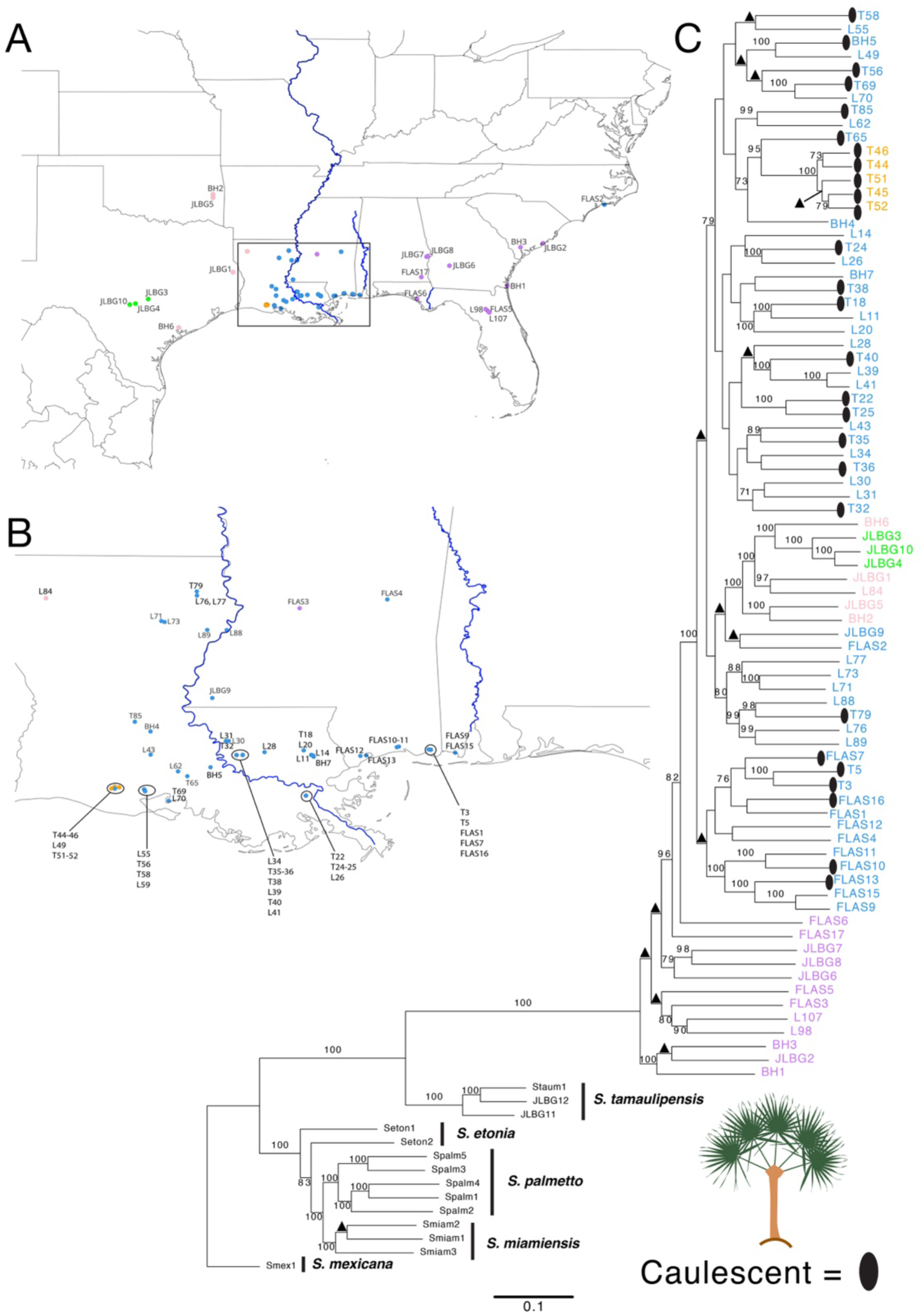
Interspecific and intraspecific relationships of Sabal minor. (A-B) Locality map showing samples (truncated “Sample Name” from Table S1) colored according to the PCA (Fig. 3). Note: Some dots representing individuals shifted on the map for stylistic purposes, for example as seen by the orange dots in (A), all the orange dots cover up the one blue dot and thus in (B) were shifted to show the presence of one individual with ancestry mostly derived from the Central Gulf Coast instead of White Lake. (C) Phylogeny of Sabal minor and other southeastern US (SE US) Sabal species inferred using RAxML. Tip labels colored following the locality map for each individual. Tip names are truncated sample names (Table S1). Black ovals designate caulescent (trunked) individuals previously known as S. louisiana. Numerical bootstrap values are shown for nodes above 70% while triangles designate nodes between 51-70%.

### Population structure and genetic differentiation

The first four PCs explained the following amount of genetic variation: 1= 5.11%, 2= 4.15%, 3= 3.45%, 4= 2.97%. Genetic clustering does not correspond to acaulescence v. caulescence stem morphology (Fig. 3A). Based on the variation of PC1, there are three genetic clusters that correspond largely to geographic locality: White Lake (orange), the Atlantic Coast (purple), and a cluster containing: the Central Gulf Coast (blue), West Gulf Coast (pink), and Central Texas (green). Along PC2, there are four clusters: White Lake, the West Gulf Coast, Central Texas, and a cluster including the Central Gulf Coast and the Atlantic Coast. With both PC1 and PC2, the PCA (Fig. 3A, 3B) shows all 78 individuals of *S. minor* cluster into roughly five clusters, the Atlantic Coast (purple), Central Gulf Coast (blue), White Lake (orange), West Gulf Coast (pink), and Central Texas (green).

**Figure 3:**
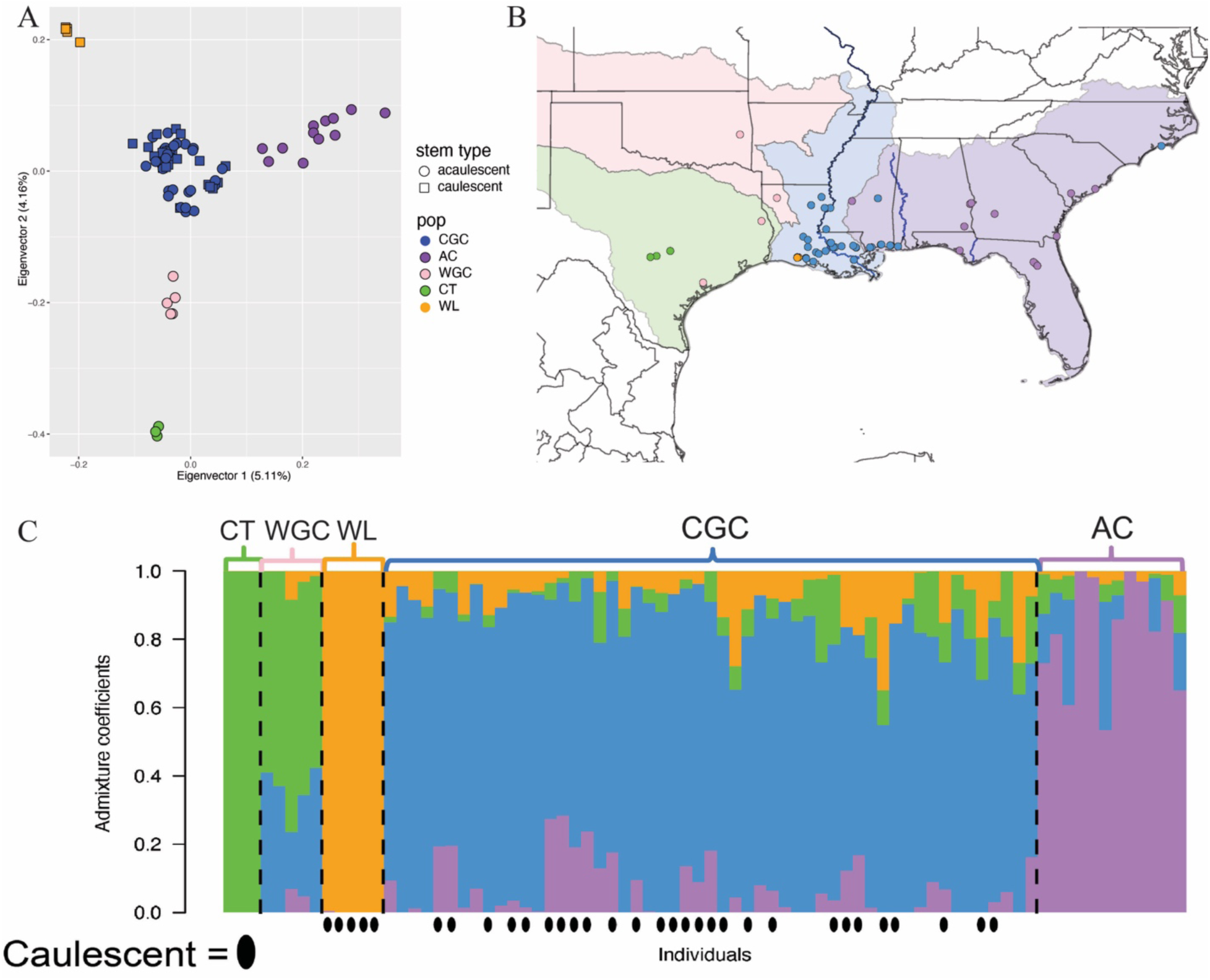
Population structure of *Sabal minor*. (A) The PCA shows the 78 individuals cluster into roughly five clusters: CGC= Central Gulf Coast, AC= Atlantic Coast, WGC= West Gulf Coast, CT= Central Texas, and WL= White Lake. (B) The five genetic clusters in the PCA correspond broadly with watershed systems in the southeastern US: South Atlantic Gulf (purple), Lower Mississippi River (blue), Arkansas-White-Red (pink), and Texas Gulf (green). The orange genetic cluster does not correspond to any watershed and is thus not regionally shaded on the map. Watershed boundaries were downloaded at the 2-digit hydrologic unit (HU2) level from the USGS 3D Hydrography Program (U.S. Geological Survey, 20250108, for each of these units: HU 03 (South Atlantic Gulf), HU 08 (Lower Mississippi), HU 11 (Arkansas-White-Red), HU 12 (Texas Gulf). (C) The most supported genetic composition for all 78 individuals parses the genotypic data into four genetic groups represented by the green, orange, blue, and purple colors. PCA population codes (CT, WGC, WL, CGC, AC) shown as dashed lines in between populations with the corresponding population names above the groups in the admixture plot. Caulescent individuals shown below the admixture plot by black ovals.

The LEA analyses indicate the best ancestral estimate is K=4 (Fig. S3). All 78 individuals of *S. minor* cluster into four populations with varying proportions of admixture (Fig. 3C). Similar to the PCA (Fig. 3A), the four populations correspond to geographic regions: White Lake, Central Texas, Central Gulf Coast, and Atlantic Coast (Fig. 3C). The West Gulf Coast does not correspond to one predominant ancestral population (Fig. 3C). The level of admixture is the highest in the Central Gulf Coast and West Gulf Coast (Fig. 3C). Corroborating the admixture results, measures of Fst indicate Central Texas and White Lake are highly differentiated from one another, with minimal gene flow between these populations, Fst= 0.5672877 (Table 1). Conversely, Fst estimates indicate the West Gulf Coast population and Central Gulf Coast are the least differentiated population pair in our study (Fst= 0.1028181).

**Table 1:** Genetic differentiation of populations. Weighted Fst measuring genetic differentiation between the five estimated populations of S. minor based on the PCA. CGC= Central Gulf Coast, AC= Atlantic Coast, WGC= West Gulf Coast, CT= Central Texas, White= White Lake.

|  | CGC | AC | WGC | CT | WHITE |
| --- | --- | --- | --- | --- | --- |
| CGC | 0 |  |  |  |  |
| AC | 0.1081736 | 0 |  |  |  |
| WGC | 0.1028181 | 0.1770169 | 0 |  |  |
| CT | 0.2200405 | 0.2814332 | 0.2383844 | 0 |  |
| WHITE | 0.2059654 | 0.3124766 | 0.39727 | 0.5672877 | 0 |

Our TreeMix analysis estimated three migration events (Fig. 4) but only one within *S. minor*, from the Central Texas to the West Gulf Coast population (Fig. 4). We did not see a trend of isolation by distance (IBD) across the entire range of *S. minor* (Table 3). However, statistically significant IBD was restricted to Central Texas and West Gulf Coast (p-value = 0.0294).

**Figure 4:**
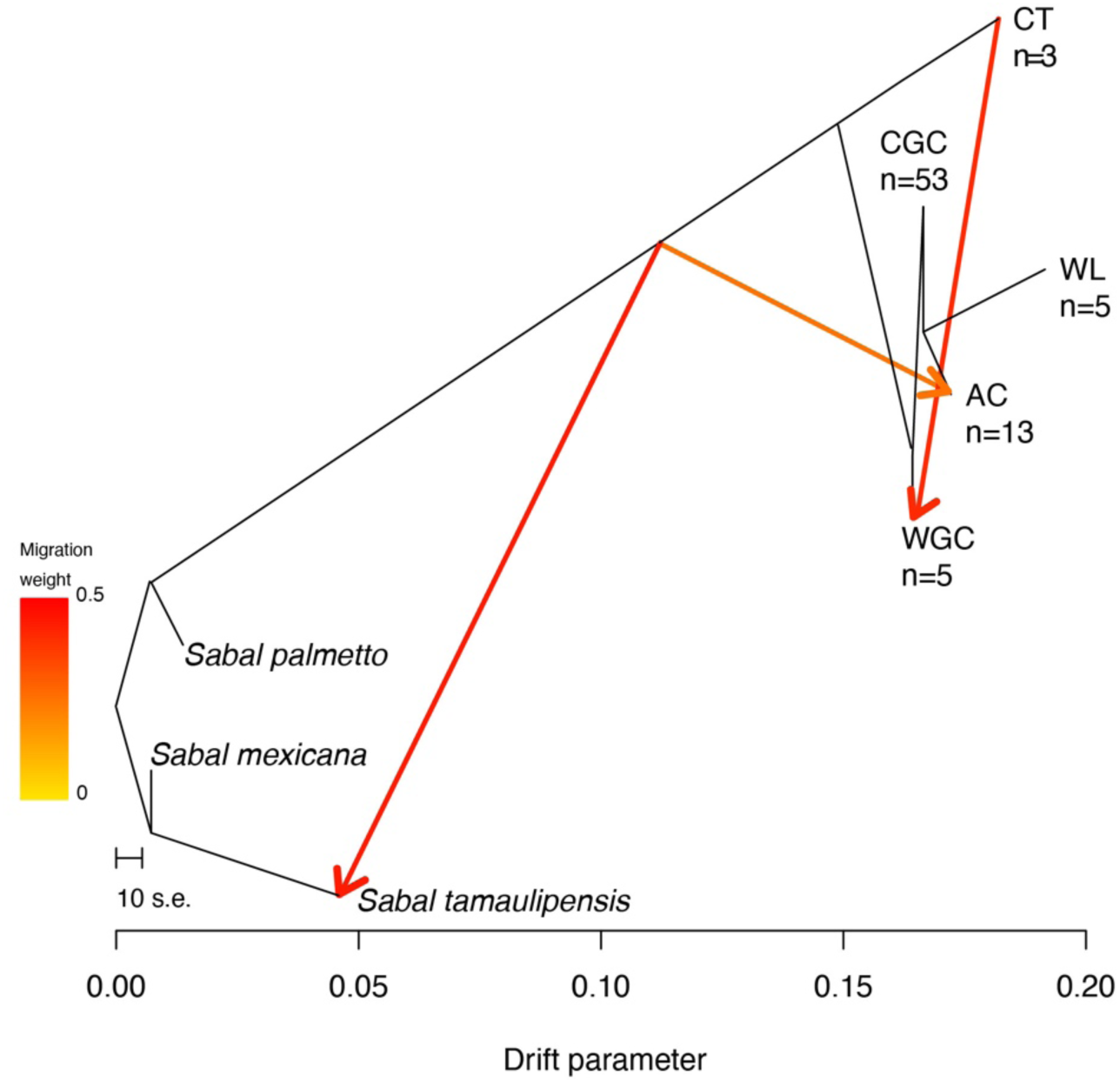
Direction of gene flow between populations. TreeMix indicates three migration events, one at the interspecific level and two within *S. minor*. Gene flow likely occurred between an ancestor of *S. minor* and *Sabal tamaulipensis.* Within *S. minor* we identified migration into the Atlantic Coast (AC) from a likely ancestral population. Lastly, TreeMix indicated gene flow from Central Texas (CT) to the West Gulf Coast (WGC). Number of individuals in each population (n).

## DISCUSSION

Our study leverages reduced representation sequence data of *S. minor* and across both stem and stemless growth forms. To facilitate SNP identification, we generated a draft genome of *S. minor*, representing one of the now four palm genomes sequenced from the sub-family Coryphoideae: *Phoenix dactylifera* (Al-Mssallem et al., 2013), *Phoenix roebelenii* (Chakravartty and Neelapu, 2023)*, Chamaerops humilis* (Bordignon, 2023), and now *Sabal minor* (present study). From population level sampling across the distribution of the species, we found that the caulescent and acaulescent forms of *S. minor* are not reciprocally monophyletic, supporting the synonymy of the two forms as one species, *S. minor* (Jacq.) Pers. Rather genetic structuring corresponds to geography where population breaks are indicated around known phylogeographic barriers, specifically the Tombigbee and Mississippi River. We also identified evidence of high genetic differentiation between two ecologically divergent populations, individuals sampled in the White Lake Conservation Area and Central Texas.

### *Sabal minor* is one polymorphic species

Our data does not indicate extensive gene flow among sampled phylogenetically and geographically proximal species of *Sabal: S. palmetto, S. etonia, S. miamiensis* and *S. minor*. This is notable because *S. minor* and *S. palmetto* are sometimes considered the putative parents of *S. x brazoriensis* (Goldman et al., 2011). Our data shows affinity of *S. x brazoriensis* with *S. minor* (Fig. S6) supporting *S. minor* as one parental contributor to this hybrid species. Additionally, our data support *S. tamaulipensis* as monophyletic and sister to *S. minor* (Fig. 2) and emphasize the original southern limit of the geographic distribution of *Sabal minor* in Texas (Bailey 1944 and Zona 1990) rather than Northeastern Mexico (Goldman 1999).

Our results also confirm that *Sabal minor* is a single species that is polymorphic for stem height, comprising both caulescent and acaulescent forms. This was anticipated given the diagnostic morphological characters for *S. minor* that are shared by both caulescent and acaulescent accessions. To this point, both forms have flattened palmate leaves, inflorescences that tend to exceed the height of the leaves, and inflorescences with long internodes that give the inflorescence a sparse appearance. Furthermore, the caulescent and acaulescent forms can co-occur (Bomhard 1943). Our genomic data support one species with the acaulescent and caulescent forms intermixed in our phylogeny (Fig. 2).

### Genetic variation is structured by geography not by caulescence

Genetic differentiation between populations of *Sabal minor* corresponds to geography rather than population structuring by stem morphotypes. Although the caulescent forms are distributed in two genetic clusters, White Lake and the Central Gulf Coast (Fig. 3), the acaulescent forms occur in all populations except White Lake indicating no genetic structuring based on phenotype. Population structure suggests four ancestral lineages that are all represented in varying admixed proportions in the Central Gulf Coast, Atlantic Coast, and West Gulf Coast subpopulations (Fig. 3). When using metrics of private allele count and nucleotide diversity (Table 2), we found the highest genetic diversity in Central Gulf Coast and Atlantic Coast. High genetic diversity in Central Gulf Coast in tandem with high phenotypic diversity is unsurprising when the ecological constraints of *S. minor* are considered. *Sabal minor* occurs across deciduous temperate forests within the southeastern United States where annual frost and snow occurs. This seasonality is considered a limitation to most palms possibly because palms are incapable of undergoing winter dormancy and they generally have only a single apical meristem (Balslev et al., 2011). *Sabal minor* may be able to tolerate frost better than most other palms in part because it is capable of maintaining its apical meristem belowground. Caulescent species of *Sabal* distributed within temperate deciduous forests of the southeastern US, such as *S. palmetto,* are limited to western Florida near Port St. Joe and Cape Fear, North Carolina (Zona, 1990), where warmer coasts provide insulation from frost. Notably, although *S. minor* largely exceeds the current geographic restrictions of other *Sabal* palms in the southeastern US, the caulescent form of *S. minor* is largely restricted to the Gulf Coast (Fig. 1A), more similar to the geographic restrictions of the other caulescent *Sabal* spp. Together, the high diversity of stem type within the Gulf Coast and high genetic diversity within both the Central Gulf Coast and Atlantic Coast groups indicates the Central Gulf Coast and Atlantic Coast as the genetic core of the species distribution of *S. minor,* consistent with the core-periphery hypothesis which suggests high genetic variation in central populations and low genetic variation in peripheral populations (Duncan et al., 2015).

**Table 2:**
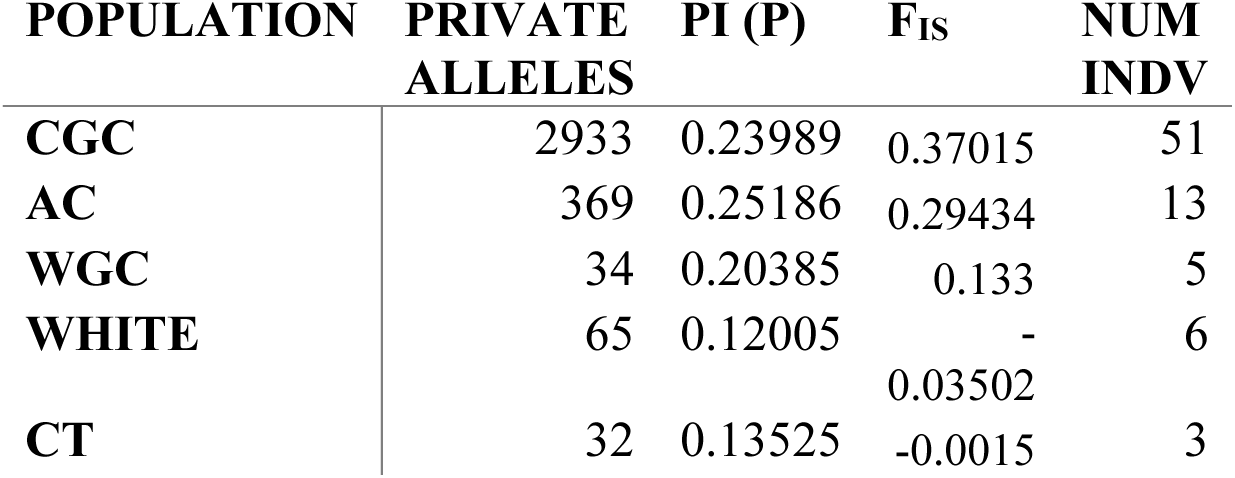
Population Summary Statistics. Statistics for each of the populations based on the PCA from variant sites only. CGC= Central Gulf Coast, AC= Atlantic Coast, WGC= West Gulf Coast, CT= Central Texas, White= White Lake.

According to nucleotide diversity (Table 2) and admixture results (Fig. 3) we observed the lowest genetic diversity in Central Texas and White Lake suggesting these populations as peripheral. Central Texas as a peripheral population is unsurprising because Central Texas is geographically isolated at the western edge of the natural range of *S. minor* (Fig. 1). Furthermore, Central Texas is located in Edward’s Plateau a semiarid ecoregion in south central Texas. The population in Central Texas is thus ecologically isolated given that other populations of *S. minor* are restricted to bottomland swamps; indeed Butler and Larson (2020) identified Edward’s Plateau as a less suitable niche than the niches occupied by the CGC and Atlantic Coast populations. The White Lake population is also ecologically divergent; this population was collected from an isolated freshwater grassland marsh within southcentral Louisiana (Fig. 3) and our data show that probably represents a founder effect, with extremely limited gene flow between White Lake and surrounding Central Gulf Coast populations.

### Phylogeographic evidence of eastern and western genetic differentiation***—***

Biota of warm temperate eastern North America are thought to have been driven into two major refugia - peninsular Florida and northeastern Mexico - during Pleistocene glacial episodes (Martin and Harrell, 1957). Key biogeographic indicators of southwestern and southeastern Pleistocene refugia and subsequent dispersals away from restricted areas include Florida-Mexico disjunct distributions and mid-continent hybridization zones resulting from post-refugial secondary contact (Martin and Harrell, 1957). Geographic breaks corresponding to Pleistocene refugia are documented in *Quercus* (Cavender-Bares et al., 2011) and *Cornus florida* (Du et al., 2024). *Sabal* appears to show this pattern by the disjunction between *S. minor* and *S. tamaulipensis* (Fig. 2). *Sabal minor* and *S. tamaulipensis* are also ecologically divergent; *S. minor* occurs in bottomland swamps and *S. tamaulipensis* occurs in submontane scrub.

Within *Sabal minor* we also observed robust evidence of eastern and western differentiated lineages, Central Texas and Atlantic Coast (Fig. 3). Notably, the Central Texas and Atlantic Coast show a similar pattern to that seen between *S. minor* and *S. tamaulipensis* where Central Texas is distributed in the semiarid Edward’s Plateau habitat while Atlantic Coast occurs within the typical bottomland swamp habitat. Phylogenetically, we observed conflicting signals regarding the phylogenetic placement of Central Texas and Atlantic Coast where RAxML (Fig. 2) and SVDquartets (Fig. S5) indicate everything diverged from Atlantic Coast while TreeMix shows Central Texas is sister to all other populations (Fig. 4). Although the topology differed between the TreeMix analyses and the other phylogenetic analyses, TreeMix suggested migration between an ancestral lineage and extant taxa, *S. tamaulipensis* and the Atlantic Coast (Fig. 4). We do not interpret this as gene flow but rather a sign that *S. minor* from the Atlantic Coast and *S. tamaulipensis* may have a higher retention of ancestral haplotypes than in other populations and species.

### Phylogeographic evidence of restricted gene flow across river systems

Rivers and river systems are well-established geographic barriers that have played dynamic roles in promoting species continuity in various biota including plants (Al-Rabab’ah and Williams, 2002; Geng et al., 2015; Nazareno et al., 2017; Sander et al., 2018), reptiles (Burbrink et al., 2000; Burbrink, 2002; Fontanella et al., 2008), and insects (Seal et al., 2015). The southeastern US is home to major river systems, such as the Mississippi River, Apalachicola River, Tombigbee River, and Red River. Phylogeographic studies have implicated these rivers, particularly the Apalachicola (Burbrink et al., 2000; Pauly et al., 2007; Walker et al., 2009; Ennen et al., 2012), the Tombigbee (Jackson and Austin, 2010; Keogh et al., 2021), and the Mississippi (Weisrock and Janzen, 2000; Near et al., 2001; Al-Rabab’ah and Williams, 2002; Saeki et al., 2011; Seal et al., 2015; Wang et al., 2023) as geographic barriers that historically sundered populations and limited gene flow. Population structure across *Sabal minor* indicates structure corresponding broadly to the Tombigbee River and to the Mississippi River delta (Fig. 3).

Perhaps it is not surprising that river systems played a role in the phylogeographic history of *S. minor* given how entwined *S. minor* is with local river systems. We identified genetically differentiated populations along the Tombigbee River, east of the river (Atlantic Coast, Fig. 3) and west of the river (Central Gulf Coast, Fig. 3). These populations both show varying degrees of admixture from all four ancestral populations (Fig. 3C). However, our results show individuals from the Atlantic Coast and Central Gulf Coast are each predominantly derived from their own ancestral populations (Fig. 3C). Atlantic Coast individuals have less admixture from the White Lake (orange) and Central Texas (green) populations than do individuals from Central Gulf Coast, which can be explained by the isolation by distance (IBD) phenomenon. Although we did not explicitly identify a trend of IBD broadly across *S. minor* and within the Central Gulf Coast and Atlantic Coast populations (Table 3), we did identify higher Fst values between the presumed source populations of the White Lake (orange) and Central Texas (green) from Atlantic Coast than from Central Gulf Coast (Table 2) to which they are geographically proximal. The lower Fst values between White Lake, Central Texas and Central Gulf Coast indirectly suggest either ongoing gene flow or relatively recent founder events leading to the establishment of the Central Texas and White Lake populations.

**Table 3:**
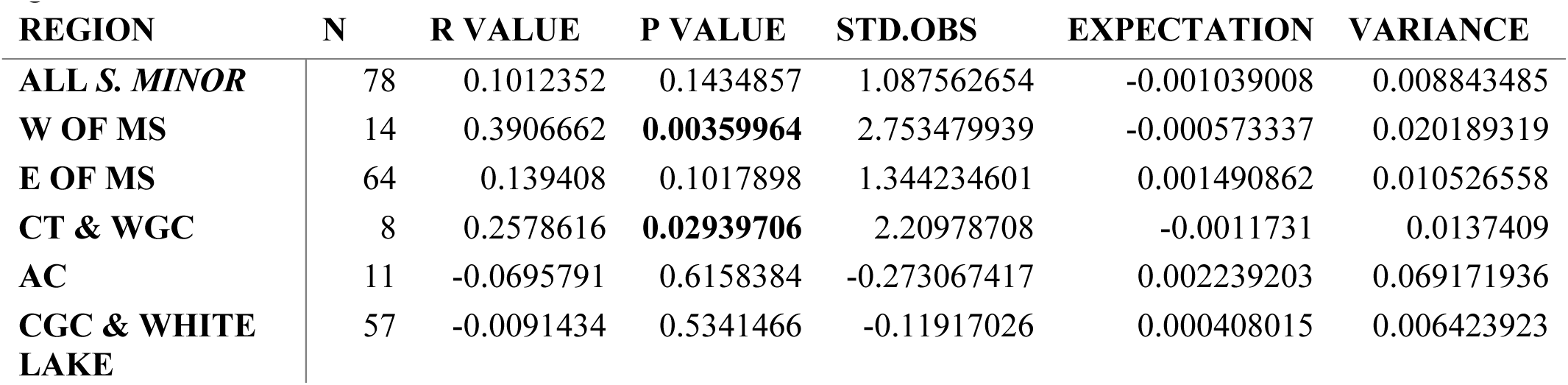
Isolation by Distance. IBD measured using variant sites for all of *S. minor*, West of the Mississippi (including Central Texas, West Gulf Coast, White Lake), East of the Mississippi (including Central Gulf Coast, Atlantic Coast) and modern river basins: Central Texas and West Gulf Coast, Atlantic Coast, Central Gulf Coast and White Lake. Bold p-values are statistically significant.

| REGION | N | R VALUE | P VALUE | STD.OBS | EXPECTATION | VARIANCE |
| --- | --- | --- | --- | --- | --- | --- |
| ALL *S. MINOR* | 78 | 0.1012352 | 0.1434857 | 1.087562654 | -0.001039008 | 0.008843485 |
| W OF MS | 14 | 0.3906662 | **0.00359964** | 2.753479939 | -0.000573337 | 0.020189319 |
| E OF MS | 64 | 0.139408 | 0.1017898 | 1.344234601 | 0.001490862 | 0.010526558 |
| CT & WGC | 8 | 0.2578616 | **0.02939706** | 2.20978708 | -0.0011731 | 0.0137409 |
| AC | 11 | -0.0695791 | 0.6158384 | -0.273067417 | 0.002239203 | 0.069171936 |
| CGC & WHITE LAKE | 57 | -0.0091434 | 0.5341466 | -0.11917026 | 0.000408015 | 0.006423923 |

The role of the Mississippi River acting as a geographic barrier is less clear from our data as Fst analyses between and among Central Gulf Coast populations indicate widespread gene flow across the Mississippi River (Fig. 3). Our analyses included dense sampling of *S. minor* individuals in central and eastern Louisiana with considerably fewer individuals sampled from western Louisiana and Texas. It is notable that within Louisiana, the populations sampled farthest west – i.e. White Lake and NWLA (L84) - are assigned to separate genetic clusters from the rest of designated Central Gulf Coast individuals (Fig. 2). This indicates the presence of some sort of barrier within central Louisiana that limited gene flow between sampled western and eastern Louisiana populations. We hypothesize that this observed break in gene flow may be the result of the dynamic nature of the Mississippi River. The Mississippi River has shifted longitudinally (Mayden, 1988; Blum et al., 2000) and has transitioned from a deep, narrow meandering course to a wide, shallow bar-braided course several times throughout its history (Rittenour et al., 2007). It is clear from our data that there is an east-west population isolation across present day Louisiana (Fig. 2), with the Mississippi River forming the largest landscape feature between the eastern (Central Gulf Coast) and western (White Lake and NWLA) populations. Genetically differentiated populations in NW Louisiana and SE Louisiana corresponding to the Mississippi River were also identified in the ringneck snake (*Diadophis punctatus*), a squamate with genetic lineages ecologically associated with floodplain forests similar to *S. minor* (Fontanella et al., 2008).

*Western populations White Lake, Central Texas, and West Gulf Coast indicate ecological differentiation (White Lake and Central Texas) and zones of introgression (West Gulf Coast)* Of the individuals sequenced from White Lake, the only one with its ancestry predominantly derived from the Central Gulf Coast was the single acaulescent individual (Fig. 3, L49). The presence of this one acaulescent individual (L49) in White Lake and the phylogenetic placement of the White Lake population deeply nested within Central Gulf Coast (Fig. 2) suggests dispersal out of Central Gulf Coast into White Lake. TreeMix did not detect the presence of gene flow between Central Gulf Coast and White Lake (Fig. 4). Notably, we detected higher genetic differentiation (as evidenced by Fst) between Central Gulf Coast and White Lake, Fst= 0.206, than between the more geographically distant Atlantic Coast and West Gulf Coast populations, Fst= 0.177 (Table 1). Given, Central Gulf Coast and White Lake are so close to one another but are more genetically differentiated from one another than Atlantic Coast and West Gulf Coast indicates a barrier to gene flow, one other than geographic distance, possibly ecologic.

The *S. minor* samples collected from White Lake in Gueydan, Louisiana were growing in an open canopy freshwater marsh (Fig. 1B, 1C) dominated by grasses and small perennial shrubs. This differs from the riparian forested habitats *S. minor* typically occurs in, where these habitats are typically subject to only seasonal flooding. The White Lake population was growing on soil that was completely inundated in December, suggesting longer periods of inundation than riparian populations experience. Furthermore, several of the White Lake palms were collected with green immature fruit in December suggesting a later flowering and fruiting time than eastern populations of *S. minor*. Because individuals predominantly derived from the White Lake ancestral population (Fig. 3C) are all caulescent, indicate some flowering time difference (between other *S. minor* populations), and occur in the different ecological setting (as compared to other *S. minor* populations), additional sampling is important to test this pattern of genetic differentiation because it hints at an interesting biological, ecological and perhaps adaptive story.

We also identified high genetic differentiation as evidenced by the admixture plot (Fig. 3C) in the Central Texas population. This population showed phylogenetic signal like White Lake where Central Texas samples were all in their own clade (Fig. 2). Also similar to White Lake, Central Texas was collected from an ecologically divergent habitat. *Sabal minor* is described as “a palm of rich soils of floodplains, levees, riverbanks, and swamps” (Zona, 1990). The Central Texas population however was collected from a semi-arid savannah habitat that occurs near the physiographic boundary between the mesic coastal plain habitat to the east and the arid great plain region to the west. Our study is limited by the number of individuals we have from Central Texas. Collection of multiple populations in Central Texas (e.g., Hill Country), north-central Texas, and along the Gulf Coast should be evaluated to test the robustness of our finding. Because, like White Lake, the genetic differentiation of the Central Texas populations could suggest adaptive processes limiting gene flow into Central Texas. Furthermore, given the similarity of habitat of the Central Texas population to *S. tamaulipensis* and the phylogenetic closeness of *S. tamaulipensis* to *S. minor*, additional investigation of the Texas populations could provide insight into its phylogeographic history. For instance, is the similar habitat between *S. tamaulipensis* and Central Texas *S. minor* an example of convergence, where both diverged from more mesic habitats? Based on morphology and/or phylogenetic placement, there are several examples within *Sabal* of transitions between mesic and xeric habitats such as *S. mexicana* and *S. guatemalensis* and *S. palmetto* to *S. etonia* (Zona, 1990). Or is the niche similarity evidence of hybridization between *S. tamaulipensis* and *S. minor*, thus the Central Texas population is of hybrid origin and inhabits the same niche as *S. tamaulipensis*? Hybrid origin of the Central Texas population from introgression between *S. mexicana* and *S. minor* was suggested by Lockett (1991). We did not infer any migration events from *S. mexicana* or *S. tamaulipensis* to the Central Texas population (Fig. 4) nor did the Central Texas individuals act as *S. x brazoriensis* (Fig. S6) where some of the Central Texas individuals were split between putative parents (*S. minor* and *S. tamaulipensis* or *S. mexicana*). Thus, hybrid origin of the Central Texas population seems the less likely explanation for the shared habitat between Central Texas and *S. tamaulipensis*. Grinage et al. (2024) found *S. minor* diverged from relatives around 12 million years ago indicating *S. minor* is ancient. It is possible the seeming divergent habitat of the Central Texas from the rest of *S. minor* could indicate *S. minor* was historically geographically and ecologically widespread. Like other palms (Baker and Couvreur, 2013), *S. minor* may have also experienced widespread extirpation (as a result of climatic cooling at higher latitudes) where Central Texas is the only surviving example of the semi-arid habitat *S. minor* also used to inhabit. Indeed, S. *minor* was identified to have one of the most sunken abaxial stomata of the entire genus, only second to a dry thorn scrub species from the Sonora (Zona, 1990). Thus, *S. minor* potentially has the physiology to make the jump from the mesic to xeric habitats. Our data suggests migration out of Central Texas to West Gulf Coast (Fig. 4) and ecological niche modelling suggests declining suitability from the Gulf Coast to Central Texas (Butler and Larson, 2020), therefore it is of conservation interest to functionally test population connectivity (seedling recruitment, pollen viability) between the Central Texas and nearby populations.

The West Gulf Coast population is admixed and not derived from a single dominant ancestral population (Fig. 3C). According to the admixture results, West Gulf Coast individuals are more genetically close to Central Texas (Fig. 3C). Furthermore, phylogenetically, the West Gulf Coast is in a clade with Central Texas samples (Fig. 2). Given this seeming genetic proximity to Central Texas, it was at first confusing that we calculated higher Fst values between West Gulf Coast and Central Texas than between West Gulf Coast and Central Gulf Coast (Table 1). Considering the geographic distribution of West Gulf Coast samples, none of these samples fit cleanly into one river system, rather they are distributed within the blue, pink, and green regions (Fig 3B). We hypothesize that these samples have strong genetic affinity to both Central Gulf Coast and Central Texas because the region represents an introgression zone. Indeed, the region was described as a phylogeographic suture zone where separated lineages of the Mississippi Embayment and the western coastal plain of southeastern Texas meet and mix (Remington, 1968). According to TreeMix, the best fit model of migration events with our data, was three migration events. The only migration event between *S. minor* populations was identified from Central Texas to West Gulf Coast suggesting that there is evidence for gene flow into the introgression zone (as defined by (Goldman et al., 2011) from Central Texas (Fig. 4). Additional to our population data showing the West Gulf Coast as an important introgression zone in *S. minor*, the only known hybrid of the genus (*S. x brazoriensis*) occurs within this zone, highlighting this geographic area as an important introgression region for *Sabal*.

## CONCLUSION

Overall, our genomic data demonstrate that the two stem growth forms of *Sabal minor* are not the result of separate evolutionary histories. Stem diversity could be the result of phenotypic plasticity or local adaptation; however, this needs to be tested by reciprocal transplant experiments. With the identified population structure in the present study, we recommend germinating seed of the caulescent type from the Central Gulf Coast and White Lake populations and transplanting in experimental plots in various sites outside of the native Central Gulf Coast and White Lake distribution to evaluate caulescence as an adaptation. Admittedly, because the caulescent form is cultivated and commercially sold, we suspect that caulescence is less likely a plastic phenotype and instead represents a locally adapted phenotype to regional environmental cues. A surprising result of our work was the high genetic differentiation identified in the White Lake and Central Texas populations. Because the White Lake and Central Texas populations may have limited gene flow into each of these populations and may inhabit geographically restricted ranges, it is of conservation interest to study these populations to inform site management plans. Lastly, our population genetic analyses show that the dynamic and long geographic history of the North American Coastal Plain has structured contemporary *S. minor* populations.

## Acknowledgments

*Herbaria*: Liberty Hyde Bailey Hortorium, University of Florida Herbarium, Shirley C. Tucker Herbarium Botanical Gardens: Montgomery Botanical Center, Juniper Level Botanic Garden *Land Managers*: Florida State Parks, Louisiana State Parks, Louisiana Wildlife Management Areas, Louisiana National Park Services: Julie Whitman

*Funding*: Cornell University (fellowships), Howard Hughes Medical Institute Gilliam Fellowship (GT15856), Montgomery Botanical Center Exploration Fund, Montgomery Botanical Center Research Fund, Cornell CALS Andrew Mellon Student Research Fund.

*Additional help from*: Lucas Majure for inspiration on the conceptualization of this project as well as silica material and Christine Bacon and Corrie Moreau for comments on the manuscript. BTI Computation Biology Center for computation resources.

## Author Contributions

ADG conceptualized this project, planned, and conducted field work, conducted the lab work, ran all the analyses except those for the genome assembly, and wrote the manuscript. JBL ran the analyses for the genome assembly, assisted with the popgen analyses, and contributed to the writing of the manuscript, as well as editing and approving the final manuscript. CDS made intellectual contributions to the conception of the project, edited, and approved the final manuscript.

## Data Availability Statement

Metadata for samples used in this study can be found in the Supplementary Materials (Table S1). Raw sequences are available on the NCBI Sequence Read Archive (SRA) accession numbers ###-####. The draft genome used to map raw reads is available for download from CoGe accession ##.

## Literature Cited

Allen, S. E. 2015. Fossil palm flowers from the eocene of the rocky mountain region with affinities to Phoenix L. (Arecaceae: Coryphoideae). International Journal of Plant Sciences 176: 586–596.

Al-Mssallem, I. S., S. N. Hu, X. W. Zhang, Q. Lin, W. F. Liu, J. Tan, X. G. Yu, et al. 2013. Genome sequence of the date palm Phoenix dactylifera L. Nature Communications 4.

Al-Rabab’ah, M. A., and C. G. Williams. 2002. Population dynamics of Pinus taeda L. based on nuclear microsatellites. Forest Ecology and Management 163: 263–271.

Austin, J. D., S. C. Lougheed, L. Neidrauer, A. A. Chek, and P. T. Boag. 2002. Cryptic lineages in a small frog: the post-glacial history of the spring peeper, Pseudacris crucifer (Anura: Hylidae). Molecular Phylogenetics and Evolution 25: 316–329.

Bagley, J. C., M. Sandel, J. Travis, M. D. L. Lozano-Vilano, and J. B. Johnson. 2013. Paleoclimatic modeling and phylogeography of least killifish, Heterandria formosa: Insights into Pleistocene expansion-contraction dynamics and evolutionary history of North American Coastal Plain freshwater biota. BMC Evolutionary Biology 13.

Bailey, L. 1934. American Palmettoes. Gentes Herbarum 3.

Bailey, L. 1944. Revision of the Palmettoes. Gentes Herbarum 6: 365–459.

Baker, W. J., and T. L. P. Couvreur. 2013. Global biogeography and diversification of palms sheds light on the evolution of tropical lineages. I. Historical biogeography. Journal of biogeography 40: 274–285.

Balslev, H., F. Kahn, B. Millan, J. C. Svenning, T. Kristiansen, F. Borchsenius, D. Pedersen, and W. L. Eiserhardt. 2011. Species Diversity and Growth Forms in Tropical American Palm Communities. Botanical Review 77: 381–425.

Bjorholm, S., J. C. Svenning, W. J. Baker, F. Skov, and H. Balslev. 2006. Historical legacies in the geographical diversity patterns of New World palm (Arecaceae) subfamilies. Botanical Journal of the Linnean Society 151: 113–125.

Blum, M. D., M. J. Guccione, D. A. Wysocki, P. C. Robnett, and E. M. Rutledge. 2000. Late Pleistocene evolution of the lower Mississippi River valley, southern Missouri to Arkansas. Geologic Society of America Bulletin 112: 221–235.

Bomhard, M. L. 1943. Distribution and character of Sabal louisiana. Journal of the Washington Academy of Sciences 33: 170–182.

Bomhard, M. L. 1935. Sabal louisiana, the correct name for the polymorphic palmetto of Louisiana. Journal of the Washington Academy of Sciences 25: 35–44.

Bordignon, A. 2023. Chromosome-scale genome assembly of the Goethe’s palm (Chamaerops humilis L.). Universita Degli Studi di Padova.

Burbrink, F. T. 2002. Phylogeographic analysis of the cornsnake (Elaphe guttata) complex as inferred from maximum likelihood and Bayesian analyses. Molecular Phylogenetics and Evolution 25: 465–476.

Burbrink, F. T., R. Lawson, and J. B. Slowinski. 2000. Mitochondrial DNA phylogeography of the polytypic North American rat snake (Elaphe obsoleta): A critique of the subspecies concept. Evolution 54: 2107–2118.

Butler, C. J., and M. Larson. 2020. Climate change winners and losers: The effects of climate change on five palm species in the Southeastern United States. Ecology and Evolution 10: 10408–10425.

Catchen, J., P. A. Hohenlohe, S. Bassham, A. Amores, and W. A. Cresko. 2013. Stacks: An analysis tool set for population genomics. Molecular Ecology 22: 3124–3140.

Cavender-Bares, J., A. Gonzalez-Rodriguez, A. Pahlich, K. Koehler, and N. Deacon. 2011. Phylogeography and climatic niche evolution in live oaks (Quercus series Virentes) from the tropics to the temperate zone. Journal of Biogeography 38: 962–981.

Chakravartty, N., and N. R. R. Neelapu. 2023. The de novo genome assembly (nuclear, chloroplast, and mitochondria) of ornamental plant pygmy date palm Phoenix roebelenii. Journal of Applied Biology and Biotechnology 11: 113–122.

Chandler, M. E. J. 1978. Supplement to the Lower Tertiary floras of southern England. Part 5. The Tertiary Research Group [ed.],. London.

Danecek, P., A. Auton, G. Abecasis, C. A. Albers, E. Banks, M. A. DePristo, R. E. Handsaker, et al. 2011. The variant call format and VCFtools. Bioinformatics 27: 2156–2158.

Darby, W. 1817. A geographical description of the state of Louisiana, the southern part of the state of Mississippi and territory of Alabama. 1st ed. James Olmstead, New York.

Darriba, Di., D. Posada, A. M. Kozlov, A. Stamatakis, B. Morel, and T. Flouri. 2020. ModelTest-NG: A New and Scalable Tool for the Selection of DNA and Protein Evolutionary Models. Molecular Biology and Evolution 37: 291–294.

Deevey, E. S. 1949. Biogeography of the Pleistocene. Bulletin of the Geological Society of America 60: 1315–1416.

Deng, T., Z. L. Nie, B. T. Drew, S. Volis, C. Kim, C. L. Xiang, J. W. Zhang, et al. 2015. Does the arcto-tertiary biogeographic hypothesis explain the disjunct distribution of Northern Hemisphere herbaceous plants? The case of Meehania (Lamiaceae). PLoS ONE 10.

Doyle, J. J., and J. L. Doyle. 1987. A rapid DNA isolation procedure for small quantities of fresh leaf tissue. Phytochemical Bulletin 19: 11–15.

Du, Z. Y., J. Cheng, and Q. Y. Xiang. 2024. RAD-seq data provide new insights into biogeography, diversity anomaly, and species delimitation in eastern Asian–North American disjunct clade Benthamidia of Cornus (Cornaceae). Journal of Systematics and Evolution 62: 1–19.

Duncan, S. I., E. J. Crespi, N. M. Mattheus, and L. J. Rissler. 2015. History matters more when explaining genetic diversity within the context of the core-periphery hypothesis. Molecular Ecology 24: 4323–4336.

Duvernell, D. D., E. Westhafer, and J. F. Schaefer. 2019. Late Pleistocene range expansion of North American topminnows accompanied by admixture and introgression. Journal of Biogeography 46: 2126–2140.

Ennen, J. R., B. R. Kreiser, C. P. Qualls, D. Gaillard, M. Aresco, R. Birkhead, T. D. Tuberville, et al. 2012. Mitochondrial DNA assessment of the phylogeography of the gopher tortoise. Journal of Fish and Wildlife Management 3: 110–122.

Fontanella, F. M., C. R. Feldman, M. E. Siddall, and F. T. Burbrink. 2008. Phylogeography of Diadophis punctatus: Extensive lineage diversity and repeated patterns of historical demography in a trans-continental snake. Molecular Phylogenetics and Evolution 46: 1049–1070.

Frichot, E., and O. François. 2015. LEA: An R package for landscape and ecological association studies. Methods in Ecology and Evolution 6: 925–929.

Geng, Q., Z. Yao, J. Yang, J. He, D. Wang, Z. Wang, and H. Liu. 2015. Effect of yangtze river on population genetic structure of the relict plant parrotia subaequalis in eastern China. Ecology and Evolution 5: 4617–4627.

Goldman, D. H. 1999a. Distribution Update: Sabal minor. Palms 43: 40–44.

Goldman, D. H. 1999b. Distribution update: Sabal minor in Mexico. PALMS-LAWRENCE- 43: 40–44.

Goldman, D. H., M. R. Klooster, M. P. Griffith, M. F. Fay, and M. W. Chase. 2011. A preliminary evaluation of the ancestry of a putative Sabal hybrid (Arecaceae: Coryphoideae), and the description of a new nothospecies, Sabal x brazoriensis. Phytotaxa 27: 8–25.

Graham, A. 1999. The Tertiary History of the Northern Temperate Element in the Northern Latin American Biota. American Journal of Botany 86: 32–38.

Greenwood, D. R., and C. K. West. 2017. A fossil coryphoid palm from the Paleocene of western Canada. Review of palaeobotany and palynology 239: 55–65.

Grinage, A. D., J. M. T. Lima, A. C. D. Maia, C. D. Specht, and L. C. Majure. 2024. Chilling Out: Cooler climates triggered divergence of Sabal (Arecaceae: Coryphoideae: Sabaleae) at the end of the Mid-Miocene Climatic Optimum. Systematic Botany 49: 567–579.

Harley, M. M. 2006. A summary of fossil records for Arecaceae. Botanical Journal of the Linnean Society 151: 39–67.

Harley, M. M., and W. J. Baker. 2001. Pollen aperture morphology in Arecaceae: Application within phylogenetic analyses, and a summary of record of palm-like pollen the fossil. Grana 40: 45–77.

Huson, D. H., and D. Bryant. 2006. Application of phylogenetic networks in evolutionary studies. Molecular Biology and Evolution 23: 254–267.

Jackson, N. D., and C. C. Austin. 2010. The combined effects of rivers and refugia generate extreme cryptic fragmentation within the common ground skink (Scincella lateralis). Evolution 64: 409–428.

Jombart, T. 2008. Adegenet: A R package for the multivariate analysis of genetic markers. Bioinformatics 24: 1403–1405.

Jones, L. N. I. I., A. D. Leaché, and F. T. Burbrink. 2023. Biogeographic barriers and historic climate shape the phylogeography and demography of the common gartersnake. Journal of Biogeography 50: 2012–2029.

Keogh, S. M., N. A. Johnson, J. D. Williams, C. R. Randklev, and A. M. Simons. 2021. Gulf Coast vicariance shapes phylogeographic history of a North American freshwater mussel species complex. Journal of Biogeography 48: 1138–1152.

De La Cerda, G. Y., J. B. Landis, E. Eifler, A. I. Hernandez, F. W. Li, J. Zhang, C. M. Tribble, et al. 2023. Balancing read length and sequencing depth: Optimizing Nanopore long-read sequencing for monocots with an emphasis on the Liliales. Applications in Plant Sciences 11.

Linck, E., and C. J. Battey. 2019. Minor allele frequency thresholds strongly affect population structure inference with genomic data sets. Molecular Ecology Resources 19: 639–647.

Mai, D. H. 1976. Fossile Früchte und Samen aus dem Mitteleozän des Geiseltales. Abhandlungen des Zentralen Geologischen Instituts 26: 93–149.

Majure, L. C., W. S. Judd, P. S. Soltis, and D. E. Soltis. 2012. Cytogeography of the Humifusa clade of Opuntia s.s. Mill. 1754 (Cactaceae, Opuntioideae, Opuntieae): Correlations with pleistocene refugia and morphological traits in a polyploid complex. Comparative Cytogenetics 6: 53–77.

Manchester, S. R. 1994. Fruits and Seeds of Middle Eocene Nut Beds Flora, Clarno Formation, Oregon. Palaeontographica Americana 58.

Manchester, S. R., T. M. Lehman, and E. A. Wheeler. 2010. Fossil Palms (Arecaceae, Coryphoideae) Associated with Juvenile Herbivorous Dinosaurs in the Upper Cretaceous Aguja Formation, Big Bend National Park, Texas. International journal of plant sciences 171: 679–689.

Martin, P. S., and B. E. Harrell. 1957. The Pleistocene History of Temperate Biotas in Mexico and Eastern United States.

Mayden, R. L. 1988. Vicariance biogeography, parsimony, and evolution in North American freshwater fishes. Syst. Zool 37: 329–355.

Morley, R. J. 2003. Interplate dispersal paths for megathermal angiosperms. Perspectives in Plant Ecology, Evolution and Systematics 6: 5–20.

Morris, A. B., C. H. Graham, D. E. Soltis, and P. S. Soltis. 2010. Reassessment of phylogeographical structure in an eastern North American tree using Monmonier’s algorithm and ecological niche modelling. Journal of Biogeography 37: 1657–1667.

Morris, A. B., S. M. Ickert-Bond, D. B. Brunson, D. E. Soltis, and P. S. Soltis. 2008. Phylogeographical structure and temporal complexity in American sweetgum (Liquidambar styraciflua; Altingiaceae). Molecular Ecology 17: 3889–3900.

Nazareno, A. G., J. B. Bemmels, C. W. Dick, and L. G. Lohmann. 2017. Minimum sample sizes for population genomics: an empirical study from an Amazonian plant species. Molecular Ecology Resources 17: 1136–1147.

Near, T. J., L. M. Page, and R. L. Mayden. 2001. Intraspecific phylogeography of Percina evides (Percidae: Etheostomatinae): An additional test of the Central Highlands pre-Pleistocene vicariance hypothesis. Molecular Ecology 10: 2235–2240.

O’Leary, S. J., J. B. Puritz, S. C. Willis, C. M. Hollenbeck, and D. S. Portnoy. 2018. These aren’t the loci you’e looking for: Principles of effective SNP filtering for molecular ecologists. Molecular Ecology 27: 3193–3206.

Ortiz, E. M. 2019. vcf2phylip v2. 0: convert a VCF matrix into several matrix formats for phylogenetic analysis. URL https://zenodo.org/records/2540861 2540861.

Pauly, G. B., O. Piskurek, and H. B. Shaffer. 2007. Phylogeographic concordance in the southeastern United States: The flatwoods salamander, Ambystoma cingulatum, as a test case. Molecular Ecology 16: 415–429.

Pickrell, J. K., and J. K. Pritchard. 2012. Inference of population splits and mixtures from genome-wide allele frequency data. Nature Precedings.

Pierce, R. L. 1961. Lower upper Cretaceous plant microfossils from Minnesota. Minnesota Geological Survey Bulletin 42: 1–86.

Prior, C. J., N. C. Layman, M. H. Koski, L. F. Galloway, and J. W. Busch. 2020. Westward range expansion from middle latitudes explains the Mississippi River discontinuity in a forest herb of eastern North America. Molecular Ecology 29: 4473–4486.

Provan, J., and K. D. Bennett. 2008. Phylogeographic insights into cryptic glacial refugia. Trends in Ecology and Evolution 23: 564–571.

Purcell, S., B. Neale, K. Todd-Brown, L. Thomas, M. A. R. Ferreira, D. Bender, J. Maller, et al. 2007. PLINK: A tool set for whole-genome association and population-based linkage analyses. American Journal of Human Genetics 81: 559–575.

Rambaut, A. 2009. FigTree (2009).

Ramp, P. F., and L. B. Thien. 1995. A Taxonomic History and Reexamination of Sabal minor in the Mississippi Valley. Principes 39: 77–83.

Reichgelt, T., C. K. West, and D. R. Greenwood. 2018. The relation between global palm distribution and climate. Scientific Reports 8.

Remington, C. L. 1968. Suture-Zones of Hybrid Interaction Between Recently Joined Biotas. Evolutionary Biology, 321–428.

Rittenour, T. M., M. D. Blum, and R. J. Goble. 2007. Fluvial evolution of the lower Mississippi River valley during the last 100 k.y. glacial cycle: Response to glaciation and sea-level change. Bulletin of the Geological Society of America 119: 586–608.

Rowan, B. A., D. K. Seymour, E. Chae, D. S. Lundberg, and D. Weigel. 2017. Methods for genotyping-by-sequencing. Methods in Molecular Biology, 221–242. Humana Press Inc.

Saeki, I., C. W. Dick, B. V. Barnes, and N. Murakami. 2011. Comparative phylogeography of red maple (Acer rubrum L.) and silver maple (Acer saccharinum L.): Impacts of habitat specialization, hybridization and glacial history. Journal of Biogeography 38: 992–1005.

Sander, N. L., F. Pérez-Zavala, C. J. Da Silva, J. C. Arruda, M. T. Pulido, M. A. A. Barelli, A. B. Rossi, et al. 2018. Rivers shape population genetic structure in Mauritia flexuosa (Arecaceae). Ecology and Evolution 8: 6589–6598.

Seal, J. N., L. Brown, C. Ontiveros, J. Thiebaud, and U. G. Mueller. 2015. Gone to Texas: phylogeography of two Trachymyrmex (Hymenoptera: Formicidae) species along the southeastern coastal plain of North America. Biological Journal of the Linnean Society 114: 689–698.

Sewell, M. M., C. R. Parks, and M. W. Chase. 1996. Intraspecific chloroplast DNA variation and biogeography of North American Liriodendron L (Magnoliaceae). Evolution 50: 1147–1154.

Small, J. 1929. Palmetto with a stem - Sabal deeringiana. Journal of the New York Botanical Garden 30: 273–284.

Small, J. K. 1926. A New Palm from the Mississippi Delta. Torreya 26: 33–35.

Soltis, D. E., A. B. Morris, J. S. McLachlan, P. S. Manos, and P. S. Soltis. 2006. Comparative phylogeography of unglaciated eastern North America. Molecular Ecology 15: 4261–4293.

Sorrie, B. A., and A. S. Weakley. 2001. Coastal Plain Vascular Plant Endemics: Phytogeographic Patterns. Castanea 66: 50–82.

Stamatakis, A. 2014. RAxML version 8: A tool for phylogenetic analysis and post-analysis of large phylogenies. Bioinformatics 30: 1312–1313.

Stephens, J. D., S. R. Santos, and D. R. Folkerts. 2011. Genetic Differentiation, Structure, and a Transition Zone among Populations of the Pitcher Plant Moth Exyra semicrocea: Implications for Conservation. PLoS ONE 6.

Sunderlin, D., J. M. Trop, B. D. Idleman, A. Brannick, J. G. White, and L. Grande. 2014. Paleoenvironment and paleoecology of a Late Paleocene high-latitude terrestrial succession, Arkose Ridge Formation at Box Canyon, southern Talkeetna Mountains, Alaska. Palaeogeography, Palaeoclimatology, Palaeoecology 401: 57–80.

Swenson, N. G., and D. J. Howard. 2005. Clustering of Contact Zones, Hybrid Zones, and Phylogeographic Breaks in North America. The American Naturalist 166: 581–591.

Swofford, D. L. 2003. PAUP*. Phylogenetic Analysis Using Parsimony. Sinauer Associates, Sunderland, Massachusetts.

Vasimuddin, M., S. Misra, H. Li, and S. Aluru. 2019. Efficient architecture-aware acceleration of BWA-MEM for multicore systems. Proceedings - 2019 IEEE 33rd International Parallel and Distributed Processing Symposium, IPDPS 2019, 314–324. Institute of Electrical and Electronics Engineers Inc.

Walker, M. J., A. K. Stockman, P. E. Marek, and J. E. Bond. 2009. Pleistocene glacial refugia across the appalachian mountains and coastal plain in the millipede genus Narceus: Evidence from population genetic, phylogeographic, and paleoclimatic data. BMC Evolutionary Biology 9.

Wang, C., Z. Yap, P. Wan, K. Chen, R. A. Folk, D. Z. Damrel, W. Barger, et al. 2023. Molecular phylogeography and historical demography of a widespread herbaceous species from eastern North America, *Podophyllum peltatum*. American Journal of Botany.

Weir, B. S., and C. C. Cockerham. 1984. Estimating F-Statistics for the Analysis of Population Structure. Evolution 38: 1358–1370.

Weisrock, D. W., and F. J. Janzen. 2000. Comparative Molecular Phylogeography of North American Softshell Turtles (Apalone): Implications for Regional and Wide-Scale Historical Evolutionary Forces. Molecular Phylogenetics and Evolution 14.

Zhang, C., S. S. Dong, J. Y. Xu, W. M. He, and T. L. Yang. 2019. PopLDdecay: A fast and effective tool for linkage disequilibrium decay analysis based on variant call format files. Bioinformatics 35: 1786–1788.

Zheng, X., D. Levine, J. Shen, S. M. Gogarten, C. Laurie, and B. S. Weir. 2012. A high-performance computing toolset for relatedness and principal component analysis of SNP data. Bioinformatics 28: 3326–3328.

Zhou, W., X. Ji, S. Obata, A. Pais, Y. Dong, R. Peet, and Q. Y. (Jenny) Xiang. 2018. Resolving relationships and phylogeographic history of the Nyssa sylvatica complex using data from RAD-seq and species distribution modeling. Molecular Phylogenetics and Evolution 126: 1–16.

Zimin, A. V., G. Marçais, D. Puiu, M. Roberts, S. L. Salzberg, and J. A. Yorke. 2013. The MaSuRCA genome assembler. Bioinformatics 29: 2669–2677.

Zona, S. 1990. A Monograph of Sabal (Arecaceae: Coryphoideae). Aliso 12: 583–666.

