## Supplemental figures for "Diversity of the dwarf palmetto: A phylogeographic analysis of the North American coastal plain and investigation of stem polymorphism in *Sabal minor* (Jacq.) Pers"

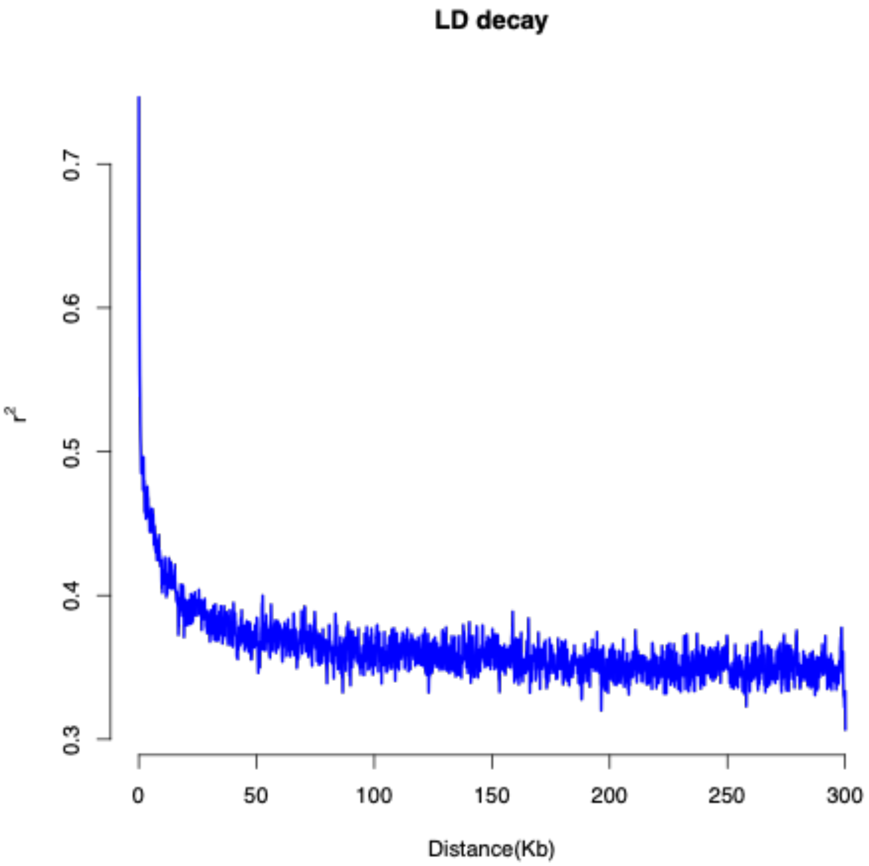


**Figure S1.** Linkage disequilibrium estimated from PopLDdecay using all 94 samples.

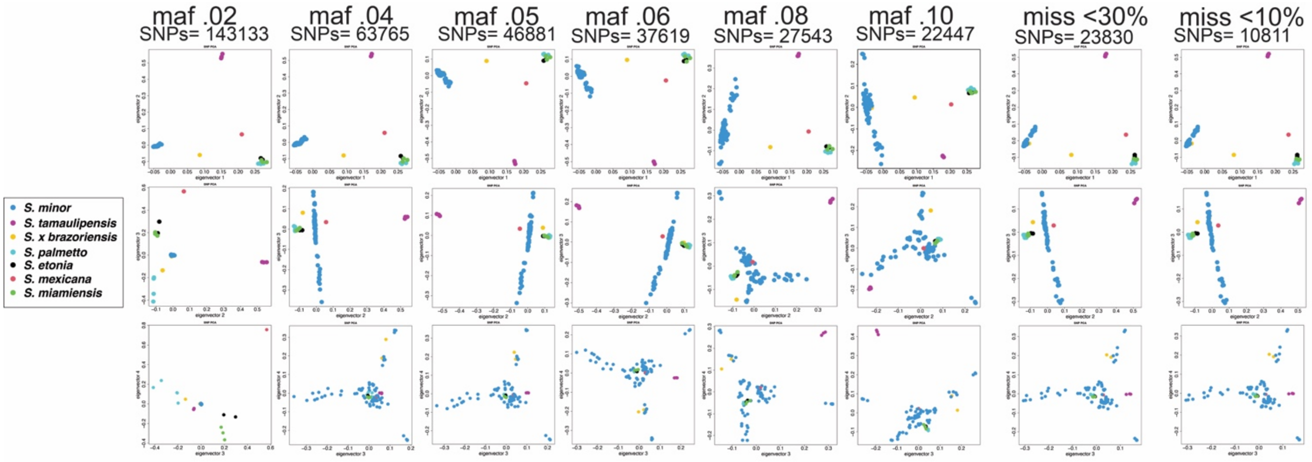


**Figure S2.** Principal component analysis panel showing robustness in filtering with varying levels of minor allele frequency (maf 0.02 to 0.1) and amount of missing data allowed (10% to 30%) in SNP filtering. Accessions are color coded according to species delimitation.

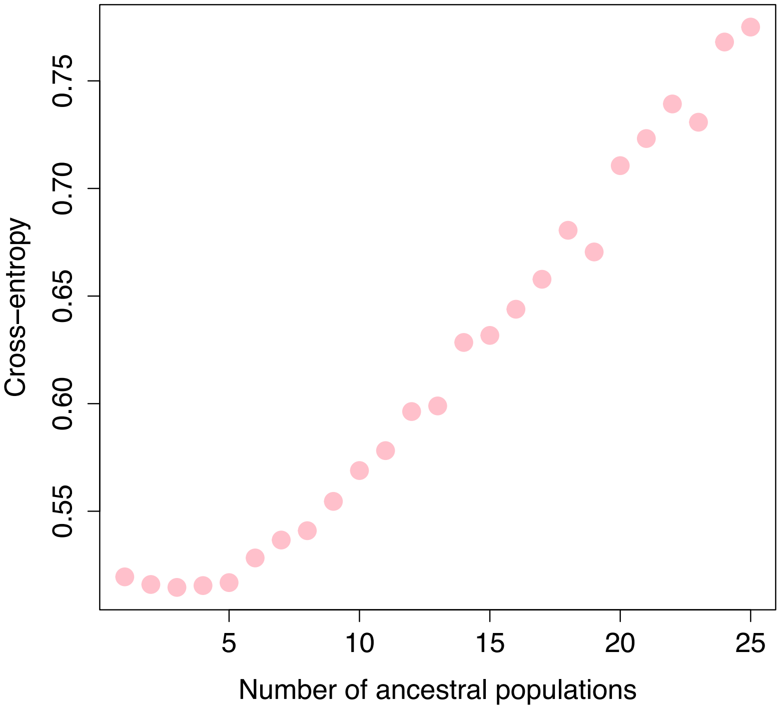


**Figure S3.** Cross-entropy plot for selection of the best number of ancestral populations from the LEA analyses.

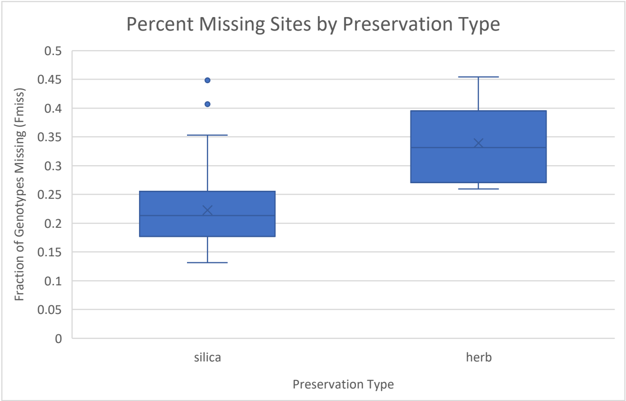


**Figure S4.** Boxplot comparing the fraction of missing SNPs between silica dried specimens (n = 85) and herbarium (n = 9) specimens for all 94 samples with the filtering threshold of missing data allowed set to 50 percent.

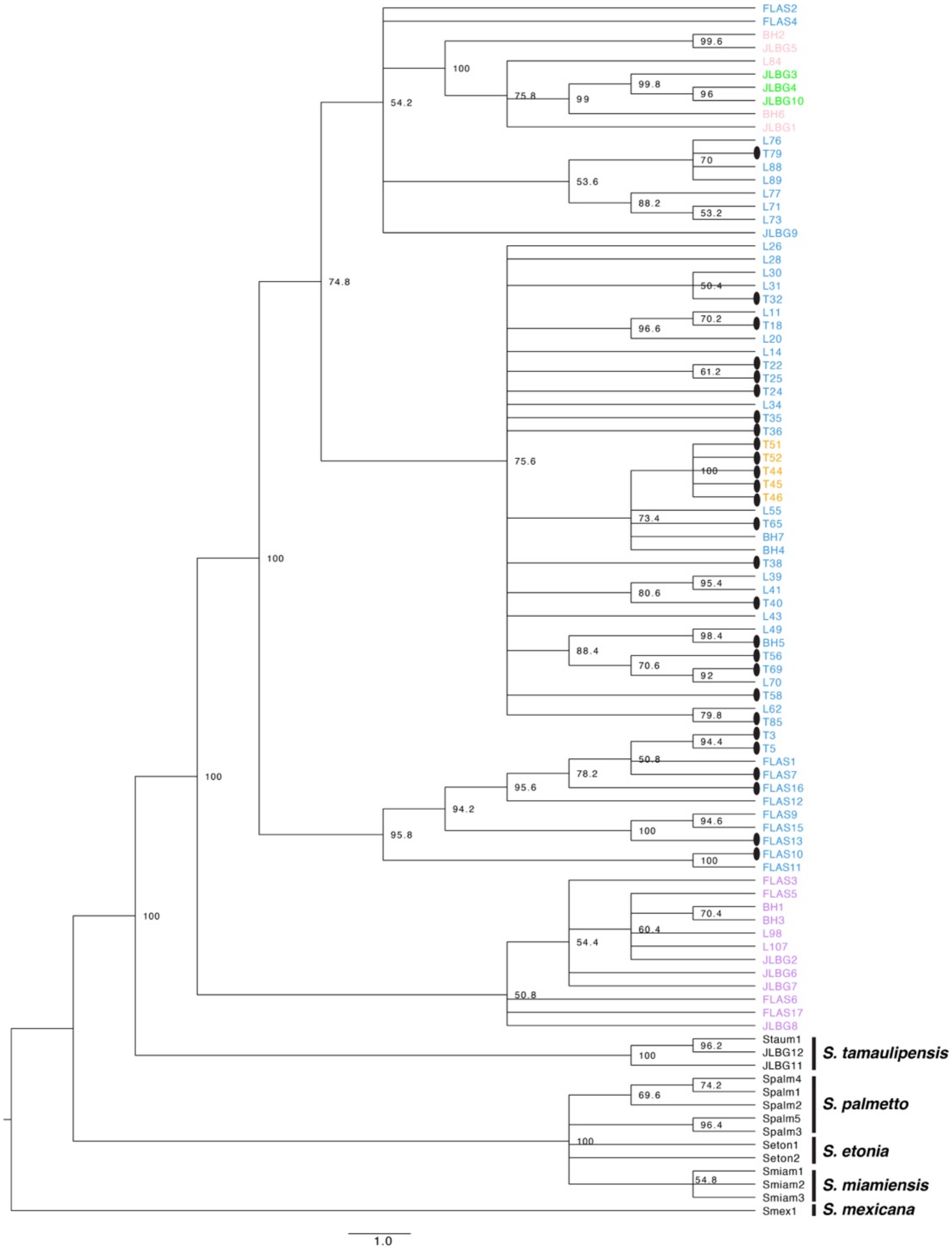


**Figure S5**. SVDQuartets majority rule consensus tree generated in PAUP. Support was assessed using 500 bootstrap replicates, only support above 50% is shown, clade support less than that is collapsed. Tips correspond to PCA population where purple= Atlantic Coast, blue= Central Gulf Coast, orange= White Lake, pink= West Gulf Coast, and green= Central Texas. Caulescent morphotype is designated by the ovals at tip terminals.

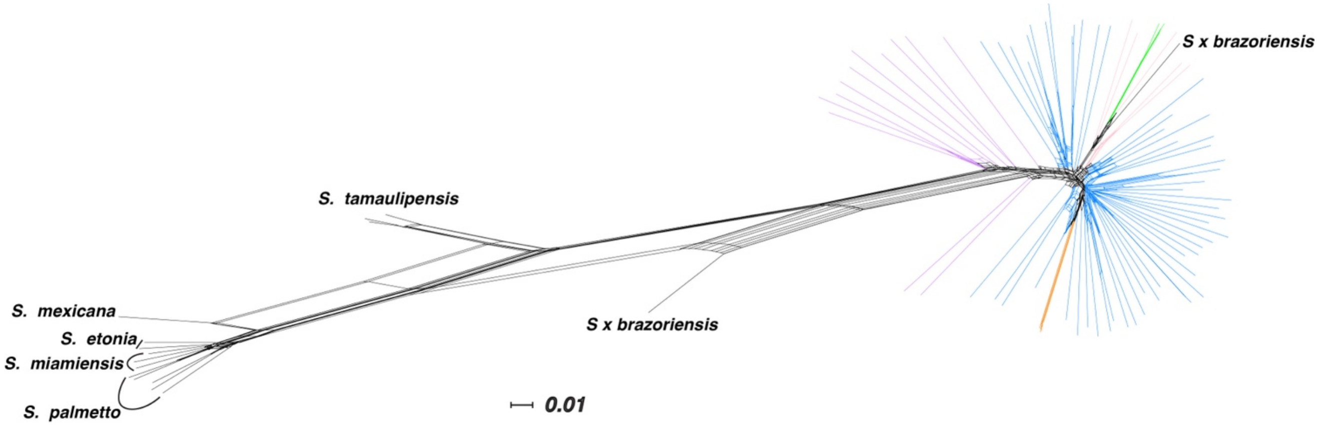


**Figure S6.** SplitsTree Neighbor network of all 94 accessions. Branches leading to terminals of *S. minor* are colored according to PCA population where purple= Atlantic Coast, blue= Central Gulf Coast, orange= White Lake, pink= West Gulf Coast, and green= Central Texas. All other species are designated by the species name.

**Appendix S1.** Additional text with of methods used for generating the genome of *Sabal minor*. (to be included in final publication)

**Table S1**. Sample metadata for all samples used in the study, including location of collections, whether they are silica or herbarium accessions, and SRA accession numbers. (to be included in final publication)
